# Conserved moiety fluxomics

**DOI:** 10.1101/2024.11.21.624666

**Authors:** Ronan M.T. Fleming, Hulda S. Haraldsdottir, German Preciat, Luojiao Huang, Ines Thiele, Amy Harms, Thomas Hankemeier

## Abstract

Fluxomics seeks to infer reaction fluxes at genome-scale by introducing isotopically labelled metabolites into a living system and measuring the enrichment of isotopic tracers in downstream metabolites. This approach has the potential to quantify fluxes at genome-scale but current approaches to infer steady-state metabolic reaction fluxes are either not tractable or not efficient at genome-scale as the corresponding computational models are high-dimensional, involve nonlinear constraints and lack mathematical guarantees of convergence to a global optimum. Herein, we introduce conserved moiety fluxomics, which is a novel, efficient, mathematically transparent, and computationally efficient method to infer metabolic reaction flux at genome-scale. We demonstrate how identification of the set of conserved moieties for a given metabolic network leads to a novel moiety graph decomposition of a stoichiometric matrix that can be used to linearly relate metabolic reaction fluxes to the rate at which conserved moieties transition between metabolites. This linear formulation avoids combinatorial explosion because the number of conserved moieties is bounded, most reactions correspond to a small number of conserved moiety transitions, it is sufficient to represent the flow of moieties that have the potential to be labelled, and it is sufficient to represent constraints from those isotopologues that are actually measured. A non-linear yet strictly convex objective is used to ensure that inferred fluxes satisfy energy conservation and the second law of thermodynamics. Numerical results demonstrate the computational tractability of conserved moiety fluxomics at genome-scale, given liquid chromatography-mass spectrometry derived stationary mass isotopologue distribution data from an in vitro dopaminergic neuronal culture fed ^13^*C* labelled glucose. Conserved moiety fluxomics unites the strengths of constraint-based modelling and metabolic flux analysis and has the potential to be further enhanced to enable inference of fluxes in whole-body metabolic models, including from non-stationary isotopic labelling data.

## 1 Introduction

Fluxomics is a range of experimental techniques [1, 2, 3, 4] to infer metabolic reaction fluxes by experimentally introducing isotopically labelled metabolite(s) into a living system and measuring the enrichment of isotopic tracers in other metabolite(s). Fluxomic techniques have the potential to quantify genome-scale fluxes *in vivo*, but there are substantial challenges to overcome. There are experimental design challenges, e.g., it is often unknown which molecular species need to be labelled to target a specific anatomical region. There are experimental implementation challenges, e.g., when non-stationary isotopic labelling is necessary to resolve particular biochemical pathways, it is experimentally challenging to sample the appropriate time course and quantify molecular species concentrations over time. The focus of this paper is to address the mathematical and computational modelling challenges of fluxomics, such as the potential for combinatorial explosion in the number of model variables or the implementation of nonlinear relationships between model variables.

Isotopomers are isomers of a metabolite that differ only in the labelling state of their individual atoms. For a metabolite containing *k* atoms, there are 2^*k*^ potential isotopomers, so the number of potential isotopomers can dramatically increase in size. This means that modelling approaches that explicitly model each potential isotopomer, e.g., [5], are impractical for application to genome-scale metabolic networks. The potential for combinatorial explosion in the number of model variables motivates the development of mathematical modelling approaches that mitigate the potential for combinatorial explosion in the number of computational model variables. An ‘elementary metabolic unit’ of a compound is ‘a moiety comprising any distinct subset of the compounds atoms’ [6]. Although, in theory, for a metabolite comprising *k* atoms, there are potentially 2^*k*^ − 1elementary metabolic units, in practice modelling a subset of elementary metabolic units is sufficient and reduced the number of model variables in the computational problem by several orders of magnitude [6], compared to explicit modelling of each isotopomer [5]. Nevertheless, estimating metabolic flux using approaches based on the concept of an elementary metabolic unit depends on the convergence of a sequence of non-linear regression problems where each iteration requires the solution of a subproblem involving a large number of coupled non-linear equations [7]. In practice, ‘it is not uncommon that algorithms get stuck in local optima’ [7], there is no established approach to test if a local optimum is also a global optimum and applications at to large networks are rare [8] are usually to relatively small networks, when compared to the size of genome-scale metabolic networks [9].

Constraint-based modelling [10] is a genome-scale modelling method that provides a molecular mechanistic framework for integrative analysis of experimental molecular systems biology data, and quantitative prediction of physicochemically and biochemically feasible phenotypic states [11]. It was established that carbon isotope labelling data can be used to constrain metabolic fluxes in a constraint-based model in a linear fashion via elementary carbon modes [12], which are analogous to elementary flux modes [13] at the level of carbon atoms. However, it it known that only subset of elementary flux modes are thermodynamically feasible [14] and the set of thermodynamically feasible fluxes is non-convex [15]. Moreover, due to the combinatorial explosion in the number of elementary carbon modes in a complete genome-scale metabolic network, this approach has not been demonstrated to be tractable at genome-scale, but rather for a metabolic network subset [12].

Constraint-based modelling has been widely applied to predict prokaryotic and eukaryotic metabolic reaction fluxes (rates). However, these predictions are usually only experimentally tested for reactions that exchange metabolites with the environment [16], or indirectly, e.g, by inferring the expression or activity of the corresponding gene product(s) [17]. As such, constraint-based modelling is currently challenged by a discordance between prediction and experimental validation. The practical requirement for established fluxomic approaches to focus on relatively small metabolic networks, contemporaneous with the growth in number of genome-scale models and modelling methods, has led to an artefactual disassociation within the metabolic network modelling community.

A computationally efficient model is one that is mathematically formulated as a problem that admits a solution obtainable with an algorithm that is computationally efficient. A computationally efficient algorithm is one with computational complexity, expressed in terms of time and space (memory) requirements, that scales polynomially with the dimension(s) of the input problem. For example, there exist computationally efficient algorithms to solve any constraint-based model that can be mathematically formulated as a convex optimisation problem [18]. Although computationally efficient algorithms allow application of many modelling approaches to genome-scale metabolic networks, no computationally efficient modelling approach has been reported that enables inference of metabolic fluxes from isotope labelling data for a complete network at genome-scale.

A conserved moiety may be descriptively defined as a group of atoms whose association within any metabolite remains invariant with respect to all reactions of a metabolic network [19]. Typically each association is represented by a chemical bonds between pairs of atoms. Every metabolite can be decomposed into a finite combination of conserved moieties. In [20] we presented an algorithmic approach to identify conserved moieties in a metabolic network by graph theoretical analysis of its corresponding atom transition network, generated from atom mapping data [21]. In [22] we refined and mathematically specified this approach and clarified an earlier conjecture that it could be used to splitting a stoichiometric matrix into the sum of a set of matrices, one corresponding to each conserved moiety.

Herein, we further refine the mathematical definition of a conserved moiety and demonstrate how identification of the set of conserved moieties for a given metabolic network leads to a novel *moiety graph decomposition* of a stoichiometric matrix that can be used to linearly relate metabolic reaction fluxes to the rate at which conserved moieties transition between metabolites. Moiety graph decomposition of a stoichiometric matrix is then applied to develop a computationally efficient constraint-based modelling method that enables inference of metabolic fluxes from isotope labelling data at genome-scale. Numerical result are presented for inference of metabolic reaction fluxes in a toy metabolic model and a genome-scale model of dopaminergic neuronal metabolism, using experimentally derived mass isotopologue distribution data from an an in vitro dopaminergic neuronal culture fed a ^13^*C* labelled glucose. Generation of mass isotopologue distribution data, atomically resolving a genome-scale metabolic model, comparison of inferred fluxes with those predicted or estimated with established approaches and biochemical interpretation of the results is the subject of a companion paper. Herein we focus on the mathematical foundations of conserved moiety fluxomics and demonstration of the computational tractability of this approach for inference of metabolic reaction fluxes at genome-scale.

## 2 Results

### 2.1 Conserved moieties

Here we describe the process to identify the conserved moieties in a biochemical network. First we start with descriptions of the input data, which are a *directed stoichiometric hypergraph*, represented by a pair of forward and reverse stoichiometric matrices, and an *atom mapping* for each reaction, then we describe the key steps in the algorithm to identify conserved moieties.

#### 2.1.1 Input data

##### Directed stoichiometric hypergraph

A metabolic network is represented by a *directed stoichiometric hypergraph* ℋ(𝒱, 𝒴(ℱ, ℛ)), which is a of oriented hypergraph, that consists of a set of vertices *m* 𝒱 ∶= {𝒱_1_, …, 𝒱_*m*_} and sequence of *n* directed hyperedges 𝒴 ∶= (𝒴_1_, …, 𝒴_*n*_). A *complex* is a multiset of molecular species. Any chemical reaction can be considered as a transformation from one complex to another. In the *j*^*th*^ reaction 𝒴_*j*_ ≔ (ℱ_*j*_, ℛ_*j*_) the tail complex is

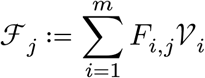

and the head complex is

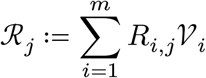

where 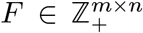 is a *forward stoichiometric matrix*, 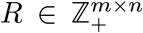 is a *reverse stoichiometric matrix*, with ℱ and ℛ being two sequences of cardinality *n*. The entry *F*_*i,j*_ is the stoichiometric number of molecule *i* consumed in the *j*^*th*^ directed reaction and the entry *R*_*i,j*_ is the stoichiometric number of molecule *i*produced in the *j*^*th*^ directed reaction. One may then define a (net) stoichiometric matrix as 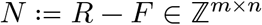

##### Atom mappings

Given a set of molecules 𝒱 and a chemical complex 𝒞(𝒱), a *complex graph* 𝒢(𝒳, 𝒴, 𝒞(𝒱)) is the disjoint union of a multiset of |𝒞| molecular graphs, where each molecular graph corresponds to one molecule 𝒱_*k*_ ∈ 𝒞. Let *n*(𝒱_*k*_) denote the atomic cardinality of molecule 𝒱_*k*_, then the total number of vertices in complex graph 𝒞 is

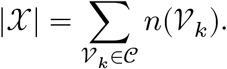

Each vertex is triply labelled with a molecular, chemical element and atomic label. The number of connected components of a complex graph is equal to the number of molecules in that complex. For example, if a complex consists of two identical molecules, that is a molecular species with stoichiometric number (multiplicity) two, then the complex graph will contain two connected components that are isomorphic up to atomic labelling.

Given a reaction ℋ ≔ {𝒫(𝒱), 𝒬(𝒱)}, an *atom transition* is a labelled edge ℰ ∶= {𝒳_*i*_, 𝒳_*j*_} that joins vertex 𝒳_*i*_ of molecule 𝒱_*k*_ in complex graph 𝒢(𝒳, 𝒴, 𝒫) with vertex 𝒳_*j*_ of molecule 𝒱_*l*_ in complex graph 𝒢(𝒳, 𝒴, 𝒬). The edge is labelled with a reaction label, which uniquely identifies a reaction. Both vertices must have the same element label, but the molecular and atomic labels may be different. That is, the element label of the vertex 𝒳_*i*_ ∈ 𝒢(𝒳, 𝒴, 𝒫) is the same as the element label of the vertex 𝒳_*k*_ ∈ 𝒢(𝒳, 𝒴, 𝒵) because an atom transition is an edge between a pair of atoms of the same element, one in each of the pair of complexes involved in a reaction. Therefore, in a reaction, the total number of atoms of each element in both complexes is the same. Given a set of molecules 𝒱 and a reaction ℋ ≔ { 𝒫(𝒱), 𝒬(𝒱)},, an *atom mapping* is a graph 𝒢(𝒳, 𝒴, ℋ{𝒫(𝒱), 𝒬(𝒱)}) formed by the disjoint union of the set of

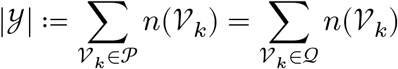

edges (atom transitions), between

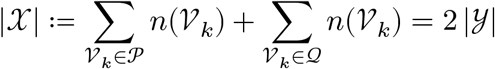

vertices (atoms). Each edge is labelled with an identical reaction label. Each vertex is labelled with an element label, a molecular label and an atomic label. Note that an atom mapping consists of |𝒴|connected components, each of which contains one edge and two vertices with identical element labels. That is, all edges of the molecular graphs of each molecule in 𝒱 are omitted. One reaction may correspond to multiple alternate atom mappings, e.g., if a molecular structure has a symmetrical subgraph, this may permit multiple alternate atom mappings that are not equivalent with respect to atomic vertex labelling but equivalent with respect to element vertex labelling. An atom mapping is required for each reaction in a metabolic network.

#### 2.1.2 Identification of conserved moieties

##### Generation of a directed atom transition multigraph

Given a directed stoichiometric hypergraph ℋ(𝒳, 𝒴{ℱ(𝒱), ℛ(𝒱)}) and an atom mapping 𝒢(𝒳, 𝒴, ℋ{𝒫(𝒱), 𝒬(𝒱)}) for each reaction, a *directed atom transition multigraph* 𝒢(𝒳, ℰ, ℋ) is a multigraph formed by the union of a set of *n* ≔ |𝒴| directed atom mappings, each of which corresponds to a reaction. The union merges vertices of atom mappings that have identical molecular, elemental and atomic labels, but duplicates edges if they have the same head and tail vertices. Each of the *p* ≔ |𝒳| vertices corresponds to an atom of an element in one of the *m* ≔ |𝒱| molecules, so each vertex is labelled with molecular, elemental and atomic labels. Each of the *t* ≔ |ℰ| edges corresponds to a directed atom transition in an atom mapping corresponding to one of the *n* ≔ |𝒴| reactions, so each edge is labelled with a reaction label. The topology of a directed atom transition multigraph is represented by an incidence matrix *T* ∈ {−1, 0, 1}^*p*×*t*^, where each row is an instance of a chemical element in a particular metabolite, and each directed edge is a directed atom transition.

A stoichiometric matrix *N* ∈ ℤ^*m*×*n*^ may be related to the incidence matrix of the corresponding directed atom transition multigraph *T* ∈ {−1, 0, 1}^*p*×*t*^ by defining two mapping matrices, as follows. Let *V* ∈ {0,1}^*m*×*p*^ denote a matrix that maps each metabolite to each atom, that is *V*_*i,j*_ = 1 if metabolite *i* contains atom j, and *V*_*i,j*_ = 0 otherwise. Each column of *V* contains a single 1 since each atom is labelled with molecular, and atomic labels and is therefore specific to a particular metabolite. Let *E* ∈ {0, 1}^*t*×*n*^ denote a matrix that maps each directed atom transition to each reaction, that is *E*_*i,j*_ = 1 if directed atom transition *i* occurs in reaction *j* and *E*_*h,j*_ = 0 otherwise. Then a stoichiometric matrix *N* can be decomposed in terms of its directed atom transition multigraph with

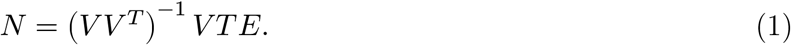

The decomposition in (1) can more easily be interpreted by rearranging terms to obtain,

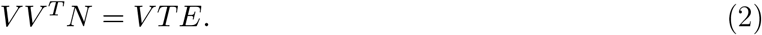

Since each column *V* of contains a single 1, the matrix 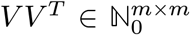 is a diagonal matrix with the total number of elements in each metabolite along the diagonal. The right hand side of (2) is therefore the internal stoichiometric matrix with each row scaled by the total number of instances of all elements in the corresponding metabolite. Every metabolite contains at least one atom so (*VV* ^*T*)^ is invertible.

##### Generation of an atom transition graph

Given a directed atom transition multigraph, an *atom transition graph* is an undirected graph 𝒜(𝒳, ℰ, ℋ) formed by removing duplicate nodes, that have identical elemental and atomic labels, and by removing edges that are identical when head and tail vertices are swapped. Each of the *p* ≔ |𝒳| vertices corresponds to an atom of an element in one of the *m* ≔ |𝒱| molecules and is labelled with molecular, elemental and atomic labels. Each of the *q* ≔ |ℰ| edges corresponds to an atom transition in one or more an atom ma*pp*ings and is unlabelled. Let *A* ∈ {−1, 0, 1}^*p*×*q*^ denote the incidence matrix of an atom transition graph. Let *E* ∈ {−1, 0, 1}^*q*×*n*^ denote a matrix that maps each atom transition to one or more reactions, that is *E*_*i,j*_ = 1 if atom transition *i* occurs with the same orientation as reaction *j, E*_*i,j*_ = −1 if atom transition *i* occurs with the opposite orientation to reaction *j* and *E h,j* = 0 otherwise. The internal stoichiometric matrix *N* can be decomposed in terms of an atom transition graph with

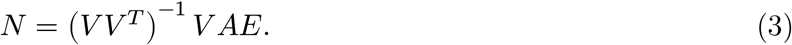

Note that, the dimension of the incidence matrices representing a directed atom transition multigraph and a corresponding atom transition graph may not be the same as the latter may have fewer columns, that is *t* ≥ *q*. Furthermore, for an atom transition graph, the matrix *E* has entries in the set {−1, 0, 1} rather than just {0, 1}, to reflect reorientation with respect to certain reactions. However, the matrix *V* is the same for the decomposition of a stoichiometric matrix in terms of a directed atom transition multigraph, or an atom transition graph.

##### Identification of the connected components of an atom transition graph

Generic graph algorithms exist to identify the connected components of a graph, and can be applied to compute the connected components of an atom transition graph. Let *C* ∈ {0, 1}^*c*×*p*^ denote a mapping between connected components and atoms in an atom transition graph, where *C*_*i,j*_ = 1 if connected component *i* contains atom *j* and *C*_*i,j*_ = 0 otherwise. This enables one to split the incidence matrix for an atom transition graph into the sum of a set of incidence matrices corresponding to its connected components.

###### Theorem 1.

(*Atom transition graph splitting*) *Let A* ∈ {−1, 0, 1}^*p*×*q*^ *be an incidence matrix for an atom transition graph* 𝒜(𝒳, ℰ, ℋ). *Let C* ∈ {0, 1}^*c*×*p*^ *be a mapping between connected components and atoms in an atom transition graph, where C*_*i,j*_ = 1 *if connected component i contains atom j and C*_*i,j*_ = 0 *otherwise, then c* = *p* − *rank*(*A*), *CA* = 0 *and the following matrix splitting exists*

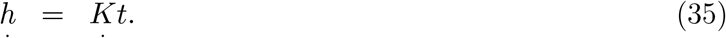

*where A*(*i*) ∈ {−1, 0, 1}^*p*×*q*^ *is an incidence matrix for the i*^*th*^ *connected component of* 𝒢(𝒳, ℰ, ℋ), *given by*

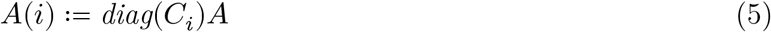

*Proof*. That *c* = *p*−rank(*A*) and *CA* = 0 are standard results from algebraic graph theory. Substituting (4) into (5), it is enough to show 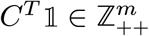 and that

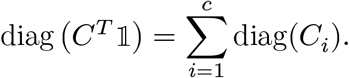

The expression on the left sums each row of *C* then places it on the diagonal of an *p* × *p* matrix. The expression on the right places each row of *C* on the diagonal of a matrix, and sums the matrices, which is equivalent to the expression on the left as the operations involved are commutative. Each entry of *C* is non-negative so *C*^*T*^ 𝟙 ≥ 0, therefore it remains to show that 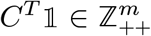. Every atom is part of one connected component, so *C*^T^ 𝟙 > 0, giving the desired result.

Note that, the connected components of an atom transition graph can be equivalently defined by mapping between connected components and atoms, or atom transitions. Next we show how connected components that are identical in particular sense may be grouped together.

##### Identification of conserved moieties via isomorphism classes of connected components

In graph theory, an isomorphism of graphs 𝒢 and ℋ is a bijection (one-to-one correspondence) between the sets of vertices of each graph such that any two vertices of 𝒢 participate in an edge if and only if the corresponding vertices in ℋ also participate in an edge. Furthermore, one can require that node labels, edge labels, or both, are preserved by a graph isomorphism. Let *A* and *B* denote the incidence matrices for two isomorphic graphs, then there exists a permutation matrix *P* such that *A* = *P* ^*T*^ *BP*. The computational problem of determining whether two finite graphs are isomorphic is called the graph isomorphism problem, and graph isomorphism algorithms exist to identify pairs of labelled graphs that are isomorphic.

Let *C* ∈ {0, 1}^*c*×*p*^ be a mapping between connected components and atoms in an atom transition graph and *C*_*i*_ denote the *i*^*th*^ row of *C*. Then *A*(*i*) ≔ diag(*C*_*i*_)*A* and *A*(*j*) ≔ diag(*C*_*j*_) are incidence matrices for the *i*^*th*^ and *j*^*th*^ connected components of the atom transition graph defined by *A*. A label preserving isomorphism between these two connected components of an atom transition graph may be represented by

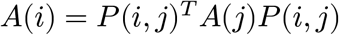

where *P*(*i, j*) ∈ {0, 1}^*p*×*q*^ is a permutation matrix representing an isomorphism between the *i* ^*ht*^ and *j*^*ht*^ connected components of an atom transition graph. If the *i*^*th*^ and *j*^*th*^ connected components of the atom transition graph are not isomorphic then *P*(*i, j*) = {0}^*p*×*q*^. When *i* = *j*, then *P*(*i, i*) ∈ {0, 1}^*p*×*q*^ is such that *A*(*i*) = *P*(*i, i*)^*T*^ *A*(*i*)*P*(*i, i*).

Given a set of graphs, a *graph isomorphism class* is a subset of graphs that are all pairwise isomorphic. Given a set of graphs, *a maximal graph isomorphism class* is subset isomorphic graphs such that no other isomorphic graph in the remainder of the set is excluded, i.e., it is of maximal cardinality. Given a set of graphs, a graph isomorphism algorithm can be applied to identify pairs of isomorphic graphs and this approach can be iteratively applied to identify each maximal graph isomorphism class. Given an atom transition graph 𝒢(𝒳, ℰ, ℋ), each connected component is a subgraph and pairs of connected components are defined to be isomorphic if their corresponding subgraphs are isomorphic, with the isomorphism preserving the molecular label of each atom. Let ℐ denote the set of maximal subgraph isomorphism classes of an atom transition graph and |ℐ| denote the number of maximal isomorphism classes. Let *D* ∈ {0, 1}^|ℐ|×*c*^ denote a mapping between isomorphism classes and connected components, where *D*_*k,i*_ = 1 if isomorphism class *k* contains connected component *i* and *D*_*k,i*_ = 0 otherwise. Then

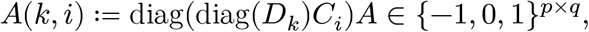

is the incidence matrix of *i*^*th*^ connected component of the atom transition matrix corresponding to the *k*^*th*^ isomorphism class ℐ(*k*). If the *j*^*th*^ connected component of the atom transition matrix is not part of the *k*^*th*^ isomorphism class then *A*(*k, j*) = {0}^*p*×*q*^.

A *conserved moiety instance* is a set of atoms with the same metabolite label, where each atom belongs to one connected component of a maximal isomorphism class of connected components of an atom transition graph 𝒜(𝒳, ℰ, ℋ). A *conserved moiety* is a species of conserved moiety instances that are identical except for their metabolite label. Each conserved moiety corresponds to one maximal isomorphism class of an atom transition graph. In a *moiety transition graph* ℳ(𝒳, ℰ, ℋ, 𝒜) each vertex is a conserved moiety instance and each edge is a *moiety transition* between a conserved moiety instance in a substrate metabolite and another conserved moiety instance, of the same conserved moiety, in a product metabolite. A moiety transition graph consists |ℐ| of connected components, each corresponding to one conserved moiety and one maximal isomorphism class of an atom transition graph.

Let *A*° (*k, i*) denote the incidence matrix of the *i*^*th*^ connected component of the atom transition graph, which also represents the canonical incidence matrix for the *k*^*th*^ isomorphism class. In a moiety transition graph, the incidence matrix of the *k*^*th*^ connected component is

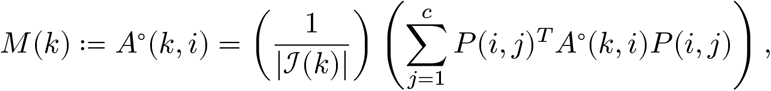

where |ℐ(*k*)| is the number of connected components of the *k*^*th*^ maximal isomorphism class ℐ(*k*) of atom transition graph 𝒜(𝒳, ℰ, ℋ). This may be interpreted as saying that *M*(*k*) is equivalent to the canonical incidence matrix for the the *k*^*th*^ maximal isomorphism class ℐ(*k*) and permutationally equivalent to each connected component in that class. Since a moiety transition graph 𝒢(𝒳, ℰ, ℋ) consists of |ℐ| connected components, the incidence matrix of a *moiety transition graph ℳ*(𝒳, ℰ, ℋ, 𝒜) is

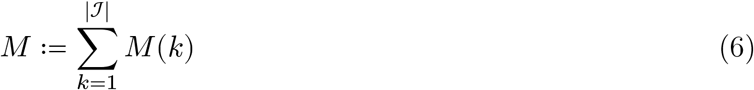

where *M*(*k*) is the incidence matrix of the *k*^*th*^ connected component, and |ℐ| is the number of maximal isomorphism classes of the corresponding atom transition graph. Henceforth, each connected component of a moiety transition graph is referred to as a *moiety subgraph*.

When an atom, or atom transition, does not participate in an isomorphic component, then the corresponding row, or column, of *P* is all zeros, respectively. It follows that *P*(*i, j*) = 0^*p*×*q*^ if the *i*^*th*^ and *j*^*th*^ connected components are not isomorphic. As defined above, the incidence matrix of a moiety transition graph has the same dimensions as the incidence matrix of an atom transition graph. However, because each conserved moiety is typically formed from more than one connected component, one can remove its zero rows and columns and define a *moiety incidence matrix M* ∈ {−1, 0, 1}^*u*×*v*^, between a set of *u* ≔ |𝒳| ≪ *p* vertices, each of which is a conserved moiety instance in a particular metabolite, and *u* ≔ |ℰ| ≪ *q* edges, each of which is a moiety transition. From algebraic graph theory, it follows that the rank of the moiety incidence matrix is equal to the number of rows minus the number of connected components of that graph, that is rank(*M*) = *u* − |ℐ|

### 2.2 Moiety splitting of a stoichiometric matrix

We aim to relate the internal stoichiometric matrix *N* ∈ ℤ^*m*×*n*^ to the incidence matrix of the corresponding moiety graph *M* ∈ {−1, 0,1}^*u*×*v*^ and its subgraphs. To this end we define two mapping matrices as follows. Let *V* ∈ {0, 1}^*m*×*u*^ denote a matrix that maps each metabolite to each moiety instance, that is *V*_*i,j*_ = 1 if metabolite *i* contains moiety instance *j*, and *V*_*i,j*_ = 0 otherwise. Each column of *V* contains a single 1 since each moiety instance is labelled with a molecular labels and is therefore specific to a particular metabolite. Let *E* ∈ {−1, 0, 1}^υ×*n*^ denote a matrix that maps each moiety transition to each reaction, that is *E*_*i,j*_ = 1 if moiety transition *i* occurs in with the same orientation in reaction *j, E*_*i,j*_ = −1 if moiety transition *i* occurs with the opposite orientation in reaction *j* and *E*_*h,j*_ = 0 otherwise. The internal stoichiometric matrix *N* can be decomposed in terms of *M, V*, and *E* as

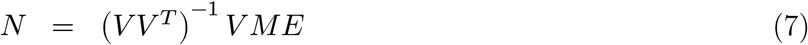

Each column of contains a single so the matrix 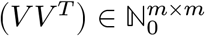 is a diagonal matrix with the total number of moiety instances in each metabolite along the diagonal. Inserting 6 into 7 one observes that

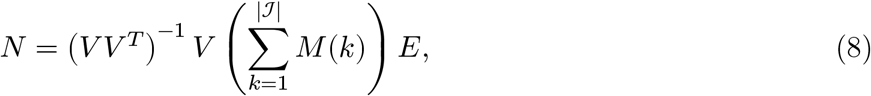

illustrating that a stoichiometric matrix, corresponding to a hypergraph, may be split into the sum of a set of moiety incidence matrices, each corresponding to a graph. The set of incidence matrices has cardinality |ℐ| = *m* − rank (*N*), that is the number of metabolites minus the number of linearly independent rows in the stoichiometric matrix.

The total number of moiety instances on the diagonal of *VV* ^*T*^ may, and typically does, consist of instances of more than one moiety. Also, every metabolite contains at least one moiety so (*VV* ^*T*^) is invertible. The decomposition in (7) may more easily be interpreted by rearranging terms to obtain

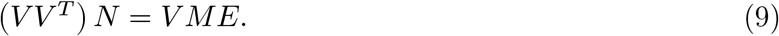

The left hand side of 9 may be interpreted as the internal stoichiometric matrix with each row scaled by the total number of instances of all moieties in the corresponding metabolite.

#### 2.2.1 Moiety decomposition of flux balance analysis

For a metabolic network, flux balance analysis [23] may be expressed as the problem

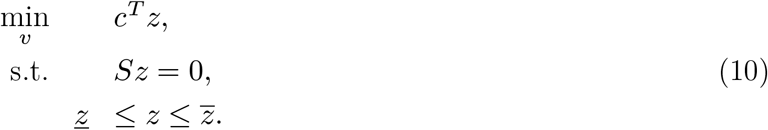

Where *<u>z</u>* ∈ ℝ^*r*^ and 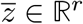 denote lower and upper bounds on net reaction fluxes, respectively. A solution *z*^⋆^ ∈ ℝ^*r*^ is a vector of steady state reaction fluxes that are optimal for some biologically motivated objective. Any stoichiometric matrix *S* ∈ ℝ may be split into subsets of columns, *S* = [*N, B*], corresponding to internal and external reactions, *N* ∈ ℤ^*m*×*n*^ and *B* ∈ ℝ^*m*×*k*^ where internal reactions are stoichiometrically consistent [24], that is 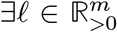 such that *N* ^*T*^ℓ = 0, and external reactions are not stoichiometrically consistent, that is 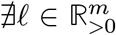 such that *B*^*T*^ ℓ = 0, as they represent net exchange of mass across the boundary of the system. In terms of internal and external stoichiometric matrices, the flux balance analysis problem can then be written as

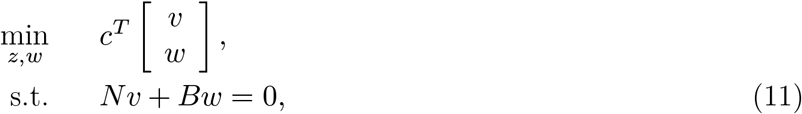

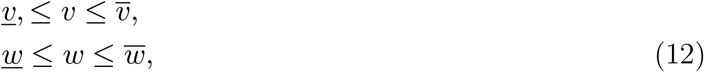

where *v* ∈ ℝ^*n*^ denotes internal fluxes and *w* ∈ ℝ^*k*^ denotes exchange fluxes. Here, *v* ∈ ℝ^*n*^ and *v* ∈ ℝ^*n*^ denote lower and upper bounds on net internal reaction fluxes, respectively, while *w* ∈ ℝ^*k*^ and *w* ∈ ℝ^*k*^ denote lower and upper bounds on net external reaction fluxes, respectively.

In general, *N* is an incidence matrix for a hypergraph with reactions corresponding to hyperedges that can connect an arbitrary number of nodes, each of which is a metabolite. Using the decomposition in (7), we can rewrite the steady state constraint in the flux balance analysis problem (11) as

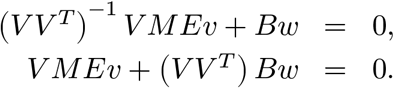

Defining a net *moiety flux* vector *y* ≔ *Ez* ∈ ℝ^υ^, one can rewrite a flux balance analysis problem as

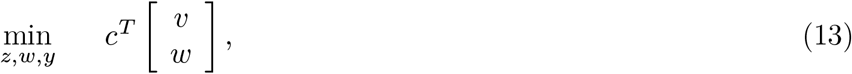

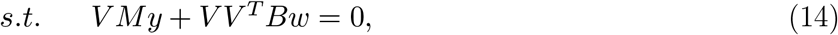

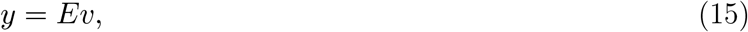

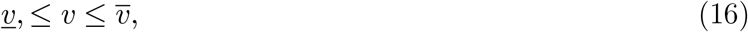

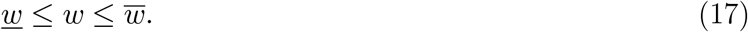

The variable *y*_*i*_ is the rate at which a moiety transitions between a substrate metabolite and product metabolite in the corresponding metabolic reaction, identified by the nonzero column of *E*_*i*_. Element *i, j* of the matrix product *VM* is the number of moieties in metabolite *i* that are part of moiety transition *j*. A moiety transition never includes more than one moiety from each metabolite. Therefore, *VM* is the incidence matrix of a graph. Writing a flux balance analysis problem in terms of moiety transitions enables us to linearly relate internal net reaction flux *v* to net moiety flux, via. *y* ≔ *Ev*

### 2.3 Steady state conserved moiety fluxomics

Here we apply moiety decomposition of a stoichiometric matrix (7), to mathematically formulate a constraint-based model that enables inference of steady state metabolic fluxes that are consistent with measured isotopologue enrichment data obtained from a system at an isotopic steady state. We formulate this model as an optimisation problem in its most general form, which assumes that each conserved moiety is isotopically labelled and that the enrichment of every isotopologue of each metabolite in the metabolic network is measured. However, these assumptions are made initially to express the generality of the approach and for clarity of mathematical exposition rather than as a necessity. After presenting this the general mathematical formulation, we explain how this approach avoids combinatorial explosion of model variables in practice.

#### Metabolite balance and thermodynamic feasibility

Steady state metabolite balance, with the option of additional constraints coupling reaction flux [25] gives rise to the constraints

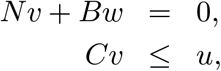

where *v* ∈ ℝ^*n*^ is net internal reaction flux, *w* ∈ ℝ^*k*^ is a vector of net external reaction fluxes, and *C* ≤ *u* represent coupling constraints. Consider the following split net internal reaction flux into unidirectional forward and reverse fluxes, *v*_*f*_ ∈ ℝ^*n*^ and *v*_*r*_ ∈ ℝ^*n*^ respectively, where *v* = *vf* − *vr*. The constraints representing steady state metabolite balance are then

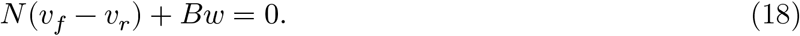

To ensure that nonzero reaction fluxes satisfy energy conservation and the second law of thermodynamics we add maximisation of the entropy of unidirectional fluxes to the objective

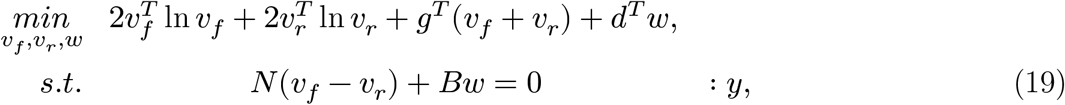

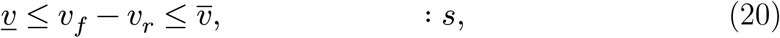

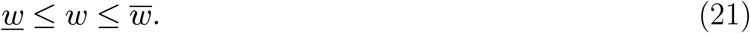

where *g* ∈ ℝ^*n*^ is a vector of free parameters that can be used to explore the entire set of thermodynamically feasible steady state fluxes [26] and *d* ∈ ℝ^*k*^ is a vector of linear objective coefficients on external net reaction fluxes. Again,*v* ∈ℝ^*n*^ and *v* ∈ ℝ^*n*^ are lower and upper bounds on net internal reaction fluxes, respectively, while respectively. *w* ∈ ℝ^*k*^ and *w* ∈ ℝ^*k*^ are lower and upper bounds on net external reaction fluxes, respectively.

With respect to unidirectional internal fluxes, the dual optimality conditions of Problem (19) are

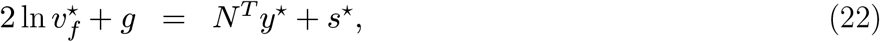

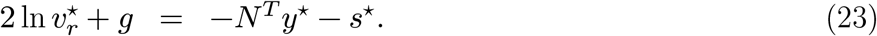

By summation of (22) and (23), one obtains

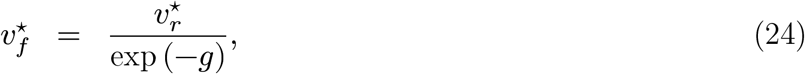

demonstrating how the free parameter *g* ∈ ℝ^*n*^ can be used to explore the steady state fluxes that correspond to different ratios of forward and reverse flux for each reaction. By subtraction of (23) from (22), one obtains

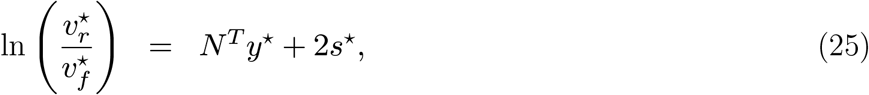

where *y*^⋆^ may interpreted as a vector proportional to the chemical potential of each metabolite and 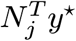 is proportional to the change of chemical potential for reaction *j*.

Let 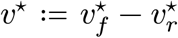. When −∞ ≤ *v*_*j*_ ≤ 0 and 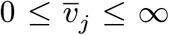 for all reactions, then the optimal dual variable 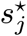 to the inequality constraints on internal reaction *j* is zero if and only if 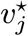 is nonzero, that is

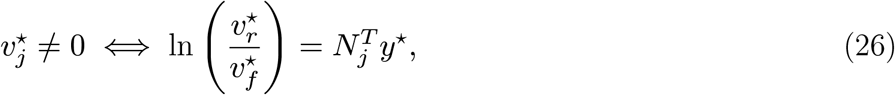

which enforces energy conservation and the second law of thermodynamics on the optimal vector of nonzero internal reaction fluxes [26]. However, when 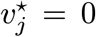.= 0, this is a relaxation of thermodynamic feasibility, since 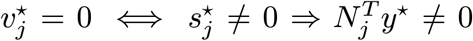.Biochemically, one may interpret this relaxation as the assumption that a zero internal reaction flux does not always imply a zero change in chemical potential. For example, a zero net flux may be consistent with a nonzero change in chemical potential when an enzyme is absent for the corresponding reaction. When *v*_*j*_ > 0 or 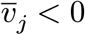 then the interpretation of 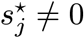 is different because a nonzero net flux may be driven in a thermodynamically infeasible direction by internal net flux bounds that force flux around stoichiometrically balanced cycles [27] independent of exchange reaction flux. Therefore, we assume −∞ ≤ *v* _*j*_≤ 0 or 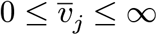.

#### Moiety balance

A key property of a conserved moiety is that it corresponds to a chemical structure that is invariant with respect to the metabolic transformations in a given network. We can use this property when modelling isotopic tracers because each conserved moiety corresponds to instances that are either isotopically labelled, or not isotopically labelled, because the chemical structure of each instance of a conserved moiety is invariant. Although it is possible to imagine multiple labelled substrates that share the same conserved moiety, where different instances of the same conserved moiety have different labelling patterns, we assume that all labelled instances of the same conserved moiety have identical labelling. That is, we assume that for each conserved moiety, if there are multiple labelled substrates sharing the same conserved moiety, they are designed so that we only need to consider either labelled conserved moiety instances or unlabelled conserved moiety instances, not multiple differentially labelled instances of the same conserved moiety. To identify mathematical terms referring to isotopic labelling, we annotate with underdot notation, that is *x* versus *x*, for example.

We assume steady state moiety balance between production, consumption and exchange with the environment, for each conserved moiety and its labelled form. We split net internal unlabelled moiety flux into unidirectional forward and reverse moiety flux vectors, *f* ∈ ℝ^υ^ and *r* ∈ ℝ^υ^ respectively, and similarly internal labelled moiety flux into unidirectional forward and reverse moiety flux vectors, *f* ∈ ℝ^υ^ and *r* ∈ ℝ^υ^ respectively. The constraints representing steady state unlabelled and labelled moiety balance are then

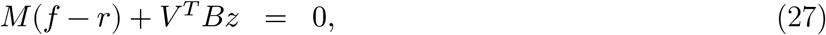

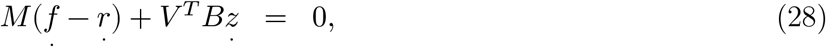

where *M* ∈ {−1, 0, 1}^*u*×υ^ is the incidencematrix of the moiety transition graph corresponding to the stoichiometric matrix in (18), *V* ∈ {0, 1}^*m×u*^ is a matrix that maps each metabolite to each moiety instance, while *z* ∈ ℝ^*k*^ and *z* ∈ ℝ^*k*^ are unlabelled and labelled net external moiety fluxes. Note that (27) is the same as (14) with the premultiplication by *V*^*T*^ omitted and net internal moiety flux split into separate forward and reverse moiety fluxes.

#### Reaction and moiety flux equivalence

Like (15), we seek to establish an equivalence between reaction flux and moiety flux, but also take into account labelling, unidirectional reactions, and the potential for differences in the direction of reactions and the corresponding moiety transitions. Let *E* ∈{−1, 0, 1}^*v*×*n*^ map moiety transitions to metabolic reactions then *E*^+^ ≔ max(0, *E*) and *E*− ≔ −min(0, *E*), so *E* = *E*^+^ −*E*^−^, then reaction and moiety flux equivalence is established by the constraints

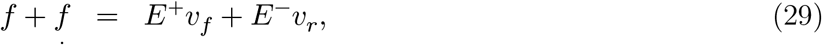

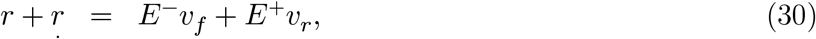

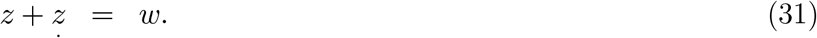

In (29) we assume that the sum of forward unlabelled flux vector and forward labelled moiety flux vectors is equal to the total reaction flux in the same direction, taking into account any differences in direction between moiety transitions and the corresponding reactions. Similarly, (30) assumes that the sum of reverse unlabelled moiety flux and labelled moiety flux vectors is equal to the total reaction flux in the same direction, taking into account any differences in directions. Equation (31) assumes that the sum of external fluxes of each unlabelled moiety, plus the corresponding labelled moiety is equal to the net external flux of the corresponding reaction. Note that the matrix that maps each metabolite to each moiety instance *V* ∈ {0, 1}^*m×u*^ has no effect on reaction direction.

#### Moiety, isotopomer and isotopologue consumption rates

We seek to relate moiety fluxes to isotopomer consumption rates. Let *F* ≔ −min(0, *M*) ∈ {0, 1}^*u*×υ^ and *R* ≔ max(0, *M*) ∈ {0, 1}^*u*×υ^ where *M* ∈ {−1, 0, 1}^*u*×υ^ is an incidence matrix for a moiety transition graph. The rates of consumption of unlabelled moieties 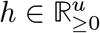 and labelled moieties 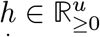 are given by

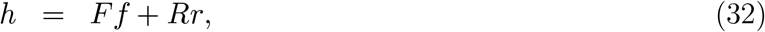

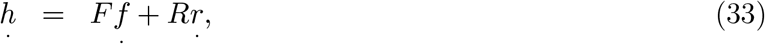

which assumes that moiety consumption by forward moiety transitions, *F f*, plus moiety consumption by reverse moiety transitions, *Rr*, equals total moiety consumption rate, and likewise for labelled moieties.

Let *K* ∈ {0, 1}^*u*×*o*^ be a matrix that maps moieties to isotopomers, where *K*_*i,j*_= 1 if the *i*^*th*^ moiety is contained in the *j*^*th*^ isotopomer, and *K*_*i,j*_ = 0 otherwise. Let *K* ∈ {0, 1}^*u*×*o*^ be a matrix that maps labelled moieties to isotopomers, where *K*_*i,j*_ = 1 if the *i*^*th*^ labelled moiety is contained in the *j*^*th*^ isotopomer, and *K*_*i,j*_ = 0 otherwise. Assuming that moiety consumption rate equals the sum of the corresponding isotopomer consumption rates *t* 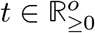,and likewise for labelled moieties, we have

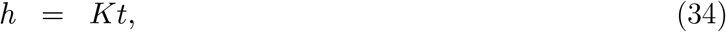

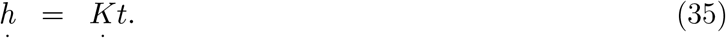

Combining (32) with (34), and combining (33) with (35), we have

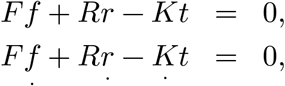

which is a linear relationship between moiety fluxes and isotopomer consumption rates.

Isotopologues are molecules that differ only in their isotopic composition. We seek to relate isotopomer consumption rates to isotopologue consumption rates 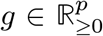 Let *Z* ∈ {0, 1}^*p*×*o*^ be a matrix that maps isotopologues to isotopomers, where *Z*_*i,j*_ = 1 if the the if the *j*^*th*^ isotopomer has the same mass as the *i*^*th*^ isotopologue and *Z*_*i,j*_ = 0 otherwise, then

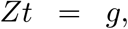

which assumes that the sum of the consumption rates of a set of isotopomers with equal mass, *Z*_*i*_*t*, equals the corresponding isotopologue consumption rate *g*_*i*_.

#### Isotopologue enrichment

We assume that isotopologue enrichment is equal to isotopologue abundance divided by the sum of the abundances of all measured isotopologues of the same metabolite. We seek to relate isotopologue consumption rate to analytical chemical measurements of isotopologue enrichment, expressed as mean and standard deviation of isotopologue enrichment for each measured isotopologue 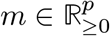 and 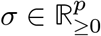 respectively. We assume that the enrichment of the *j*^*th*^ isotopologue *mj* is equal to the rate of consumption of the *jth* isotopologue *gj*, divided by the sum of the rates of isotopologue consumption of the set of isotopologues of the same metabolite as the *j*^*th*^ isotopologue Φ(*j*) that is

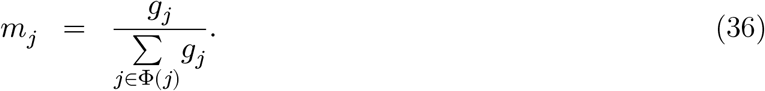

However, rather than imposing this constraint directly, we introduce a residual variable *s*_*j*_ then seek to minimise the square of this residual, weighted by the inverse of the standard deviation of isotopologue enrichment, that is

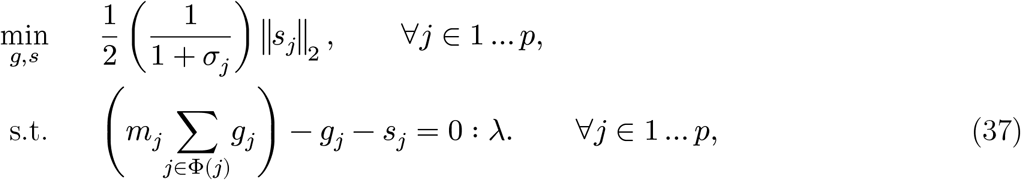

From the optimality conditions, one obtains

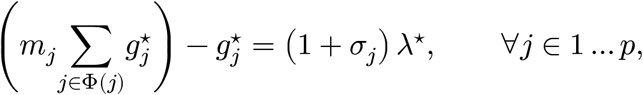

so any deviation from the hard optimisation constraint in (36) is penalised less if the standard deviation of isotopologue enrichment σ_*j*_ is large. Problem (37) is quadratic optimisation subject to linear constraints of the form

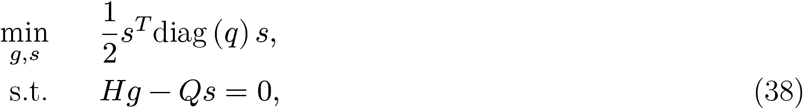

where 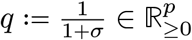 and *H* ∈ ℝ^*p*×*p*^ and *Q* ∈ ℝ^*p*×*p*^ are matrices populated with isotopologue enrichment data in the appropriate sparsity pattern to implement the constraints in Problem (37).

#### 2.3.1 Mathematical formulation of conserved moiety fluxomics

Combining the aforementioned set of objectives and constraints, we obtain the following convex optimisation problem

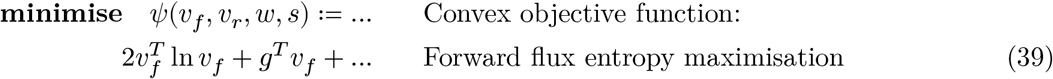

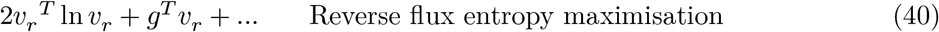

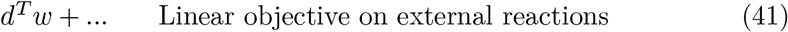

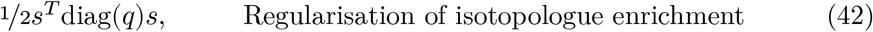

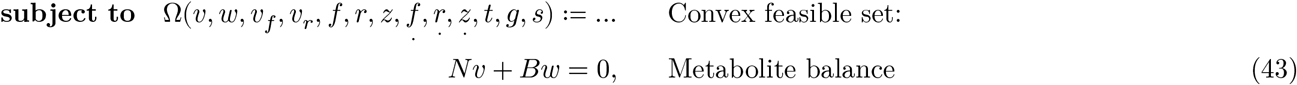

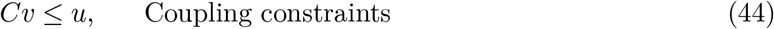

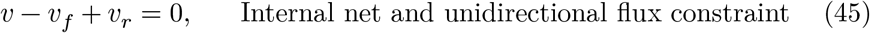

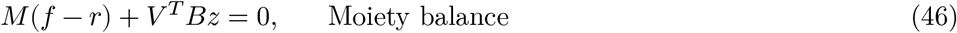

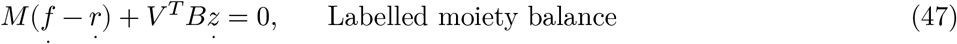

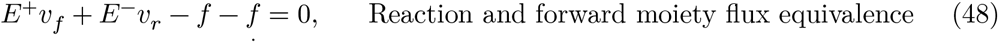

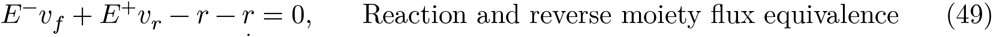

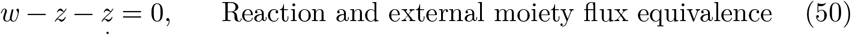

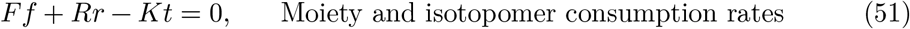

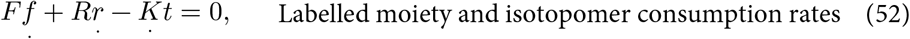

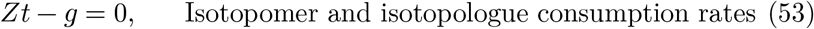

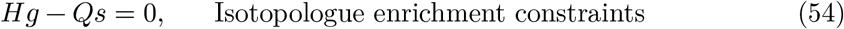

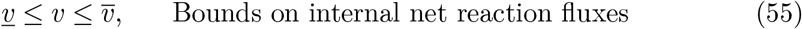

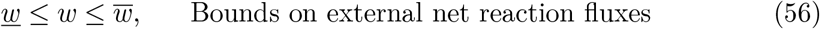

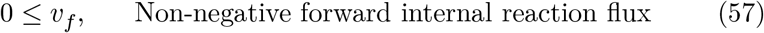

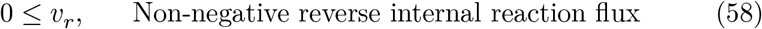

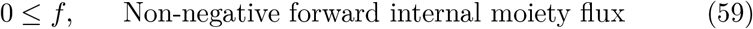

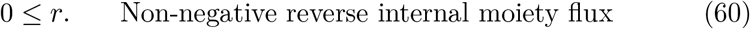

The variables are internal net flux *v* ∈ ℝ^*n*^, internal forward reaction fluxes *vf* ∈ ℝ^*n*^, internal reverse reaction fluxes *v*_*r*_ ∈ ℝ^*n*^, external net reaction fluxes *w* ∈ ℝ^*k*^, internal forward moiety fluxes *f* ∈ ℝ^υ^, internal reverse moiety fluxes *r* ∈ ℝ^υ^, external net moiety fluxes *x* ∈ ℝ^*k*^, internal forward labeled moiety fluxes *f* ∈ ℝ^υ^, internal reverse labelled moiety fluxes *r* ∈ ℝ^υ^, external net labelled moiety fluxes *x* ∈ ℝ^*k*^, isotopomer consumption rates 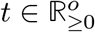,isotopologue consumption rates 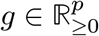 and isotopogue enrichment constraint residual *s* ∈ ℝ^*p*^.

The data are parameters to explore the set of thermodynamically feasible steady state fluxes *c* ∈ ℝ^*n*^, linear objective coefficients on external net reaction fluxes *d* ∈ ℝ^*k*^, quadratic penalties on violation of isotopologue enrichment constraints *q* 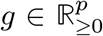, an internal stoichiometric matrix *N* ∈ ℤ^*m*×*n*^, an external stoichiometric matrix *B* ∈ ℝ^*m*×*k*^, a moiety transition graph incidence matrix *M* ∈ {−1, 0, 1}^*u*×*v*^ where *F* ≔ −min(0, *M*) ∈ {0, 1}^*u*×*v*^ and *R* ≔ max(0, *M*) ∈ {0, 1}^*u*×*v*^, a matrix that maps each metabolite to each moiety instance *V* ∈ {0, 1}^*m*×*u*^, a matrix that maps moiety transitions to metabolic reactions *E* ∈ {−1, 0, 1}^*v*×*n*^ where *E*^+^ ≔ max(0, *E*) and *E*^−^ ≔ −min(0, *E*)), a matrix that maps moieties to isotopomers *K* ∈ {0, 1}^*u*×*o*^, a matrix that maps labelled moieties to isotopomers *K* ∈ {0, 1}^*u*×*o*^. a matrix that maps isotopomers to isotopologues *Z* ∈ {0, 1}^*p*×*o*^, and two matrices populated with isotopologue enrichment data *H* ∈ ℝ^*p*×*p*^ and *Q* ∈ ℝ^*p*×*p*^.

#### 2.3.2 Combinatorial implosion

The mathematical formulation of conserved moiety fluxomics in (60) is a minimisation of a nonlinear convex function *ψ* over a polyhedral convex set of the form Ω ≔ {*x* ∣ *Ax* ≤ *b*}. There exist computationally efficient algorithms to solve convex optimisation problems [18], however the computational complexity is calculated as a function of the problem dimensions. Computationally, one still requires that the problem dimension is not too large as to make the problem intractable. Problem (60) has the potential to combinatorially explode if each conserved moiety is isotopically labelled and that enrichment of every isotopologue of each metabolite in the metabolic network is measured. However, in practice, this is never the case and therefore one can reduce the size of Problem (60) dramatically.

A moiety transition graph 𝒢(𝒳, ℰ, ℋ) consists of a maximum of |ℐ| = *m* − rank(*N*) connected components, where *m* is the number of metabolites in the metabolic network given by the stoichiometric connected matrix *N*. If each metabolite contained one instance of each moiety, a moiety transition graph could *m* × |ℐ| vertices, but that is improbable as in a typical genome-scale metabolic network, few metabolites have many moiety instances while most metabolites have few moiety instances (Figure 1). If each reaction contained one moiety transition of each moiety, a moiety transition graph could consist of *n* × |ℐ| vertices, but that is improbable as in a typical genome-scale metabolic network, few reactions have many moiety transitions while most reactions have few moiety transitions (Figure 1).

**Figure 1:**
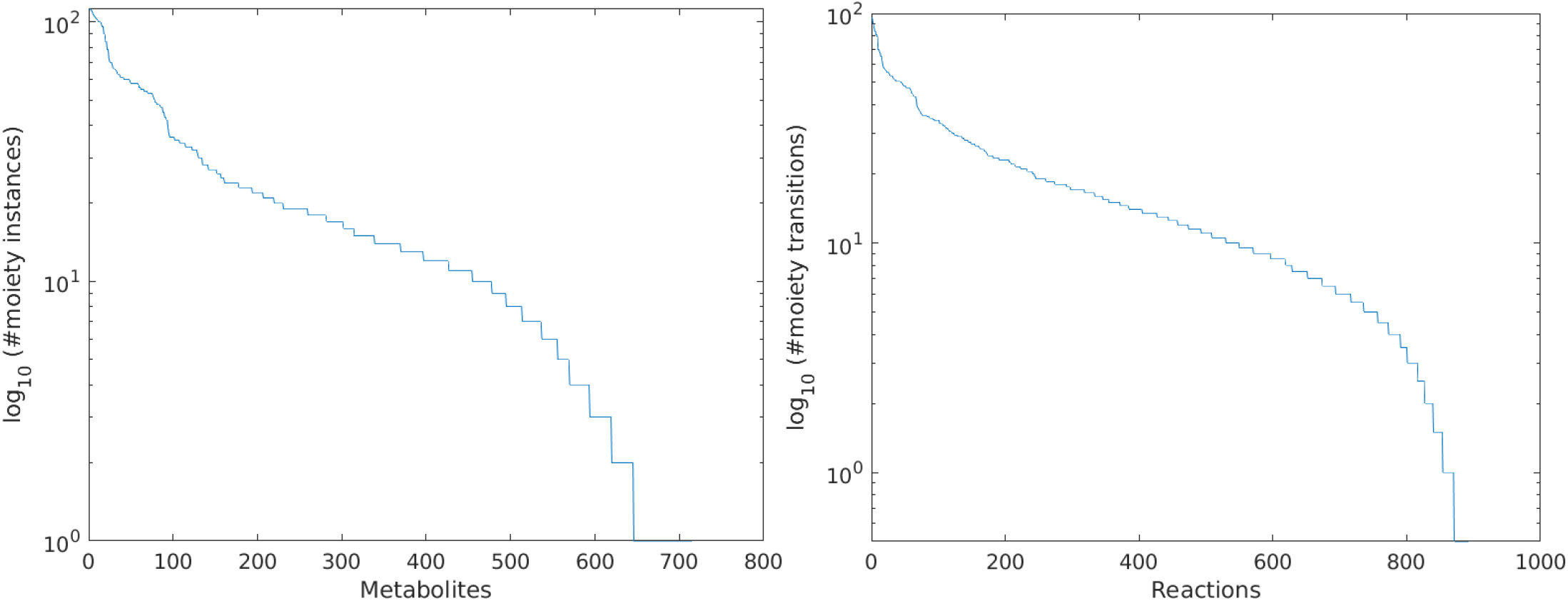
Enumeration of moiety instances per metabolite (left) and moiety transitions per reaction (right) in a genome-scale metabolic model of dopaminergic neuronal metabolism [16].

**Figure 2:**
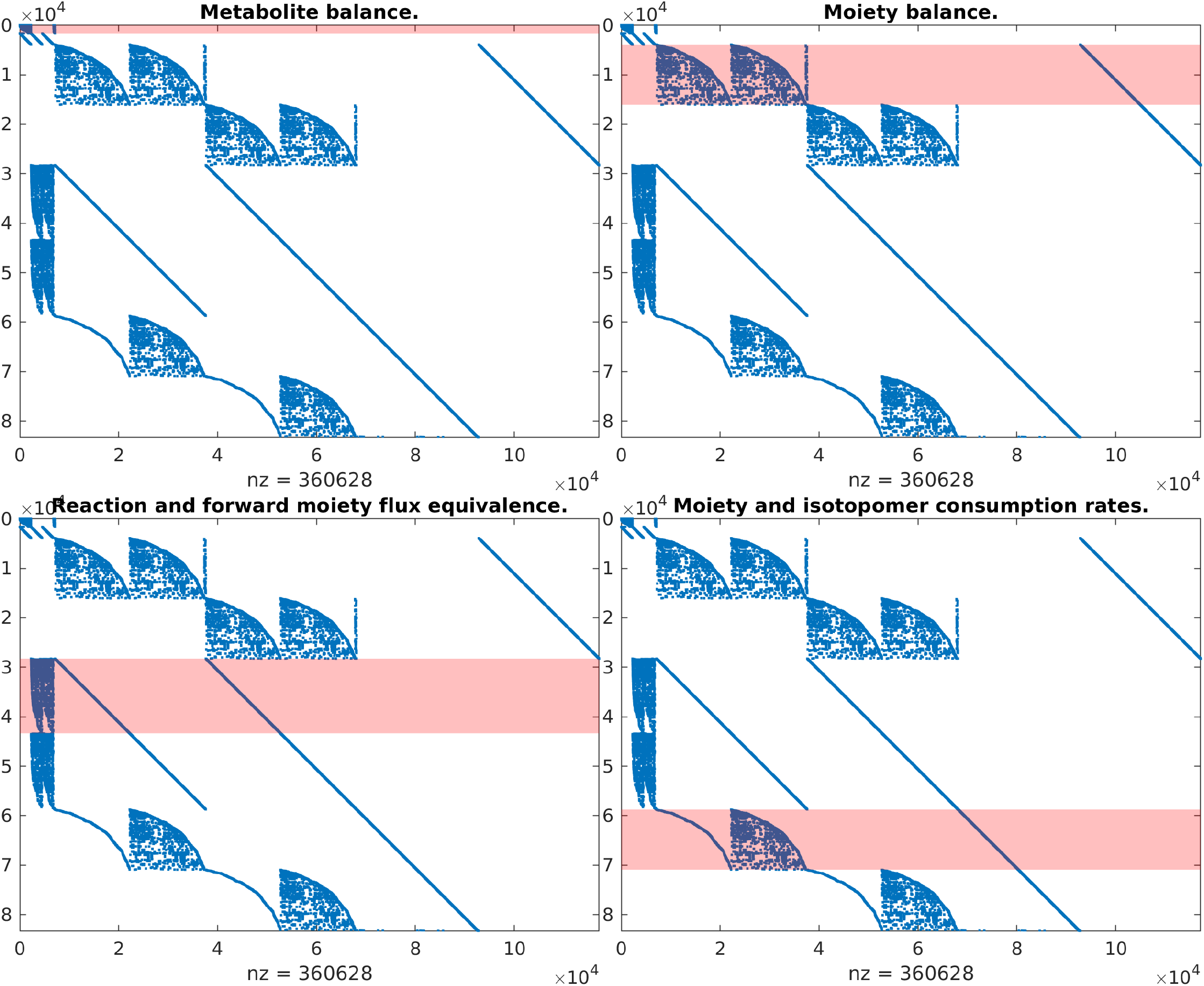
Each panel is the same moiety constraint matrix for the genome-scale model of dopaminergic neuronal metabolism, *A* ∈ ℝ^81,015×114,880^ in Table 1, where rank (*A*) = 83, 165. In the moiety constraint matrix, the rows are in the same order as the constraints in Problem (60) and the columns are in the same order as variables in in Problem (60). Highlighted metabolite balance constraints (*N v* + *Bw* = 0, top left), moiety balance constraints (*M*(*f* − *r*) + *V* ^*T*^ *Bz* = 0,top right), forward reaction and moiety flux equivalence (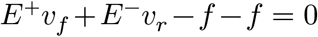, bottom left), and constraints relating moiety and isotopomer consumption rates (*F f* + *Rr* − *Kt* = 0, bottom right). The corresponding labelled versions are below each of the latter latter three constraints.

In any case, when applying Problem (60) in practice, it is sufficient to only represent the moiety subgraphs corresponding to the labelled moieties within the typically small set of labelled substrate metabolites, which are designed at the outset of the experiment. This dramatically reduces the size of the moiety transition graph incidence matrix. That is 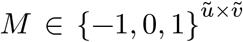 where 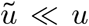 and 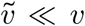. Consequently, it also dramatically reduces the size of 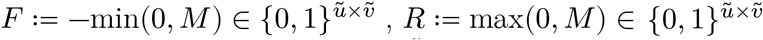 the matrix that maps each metabolite to each moiety instance 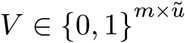, the matrix that maps moiety transitions to metabolic reactions 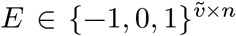, the matrix that maps moieties to isotopomers 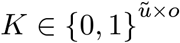,and the matrix that maps labelled moieties to isotopomers 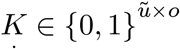.

For a metabolite containing *k* moiety instances, assuming each moiety instance is either unlabelled or labelled with the same pattern, there are 2^*k*^ potential isotopomers, so the number of potential isotopomers has the potential to dramatically increase in size. Even though there are always less isotopologues than isotopomers, any combinatorial explosion in the number of isotopomers also has the potential to increase the number of isotopologues. However, when applying Problem (60) in practice, it is sufficient to only represent those isotopologues that are actually measured. Unmeasured isotopologues are unconstrained, so do not add to the constraints on Problem (60). This dramatically reduces the size of the two matrices populated with isotopologue enrichment data 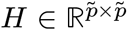 and 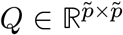, where 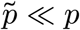, the matrix that maps isotopomers to isotopologues 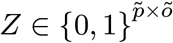, where 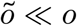, the matrix that maps moieties to isotopomers 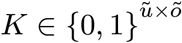,and the matrix that maps labelled moieties to isotopomers 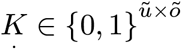.

Fundamentally, by modelling isotope labelling as a combination of a bounded number of irreducible metabolic entities, i.e. conserved moiety instances, hard coupled to network fluxes and only implementing constraints on moiety transition fluxes arising from labelled substrates and measured isotopologues, means that the conserved moiety fluxomic model formulated as Problem (60) permits a combinatorial implosion of what has the potential to become a combinatorial explosion when unnecessary variables are modelled explicitly.

### 2.4 Numerical experiments

Table 1 summarises the numerical results of applying conserved moiety fluxomics to a toy model and to tracer-based metabolomic data from *in vitro* human dopaminergic neuronal cell culture that was fed with an isotopically labelled substrate, as described in detail in a companion paper (Huang et al, unpublished). Briefly, fully ^13^ labelled glucose was fed to a dopaminergic neuronal culture, obtained by differentiation of neuroepithelial stem cells, derived from induced pluripotent stem cells [28]. Mass isotopologue distributions were obtained for 10 metabolites (52 isotopologues) by liquid chromatography-mass spectrometry of labelled intracellular samples, as described elsewhere^1^. Given an established genome-scale model of dopaminergic neuronal metabolism (`iDopaNeuroC`, [16]), 93% (2,103/2,270) of its internal reactions were algorithmically atom mapped using Reaction Decoder Tool [29], which was previously established as highly accurate when compared to manual atom mapping of biochemical reactions, in an independent comparative evaluation of atom mapping algorithms for balanced metabolic reactions [21].

**Table 1:**
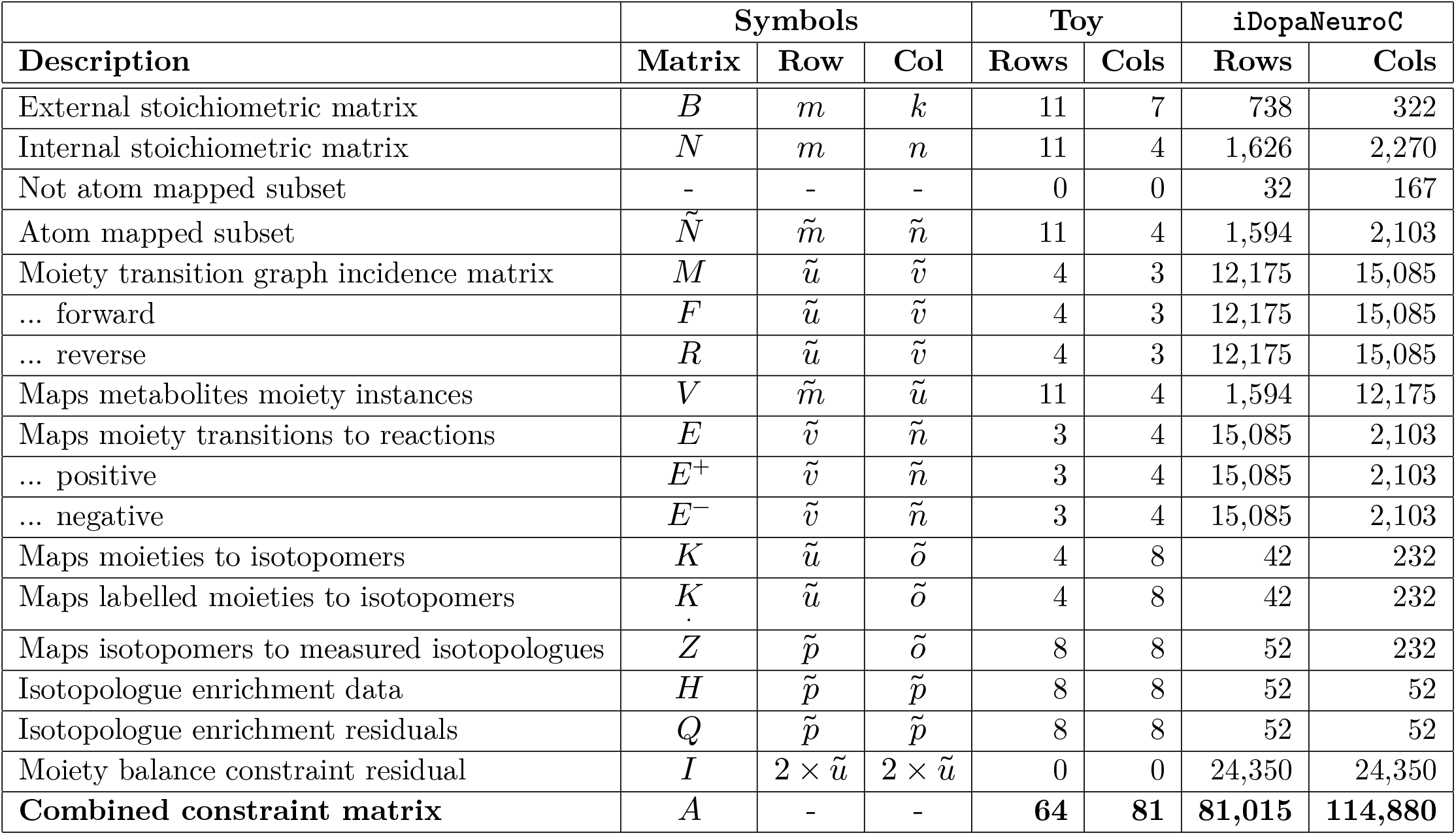
Dimensions of matrices in Problem (60) when conserved moiety fluxomics is applied to a small network (toy) and a genome-scale model of dopaminergic neuronal metabolism (iDopaNeuroC, [16]). All matrices are sparse. The atom mapped subset of the internal stoichiometric matrix corresponding 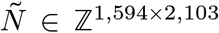 has rank 1,451, so in principle the number of conserved moieties should be 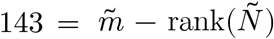 however |ℐ| = 195 conserved moieties were identified, corresponding to 136 linearly independent moiety vectors, that is rank (*L*) = 136. Since 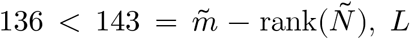 is not a complete basis for the left nullspace of 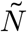.

Given a directed stoichiometric hypergraph of the `iDopaNeuroC` model, represented by an internal stoichiometric matrix with 1,594 rows (metabolites) and 2,103 columns (reactions), and an atom mapping for each reaction, we identified the conserved moieties of the atom mapped subset of the `iDopaNeuroC` model, as described in Subsection 2.1. Subsequently, as described in Subsection 2.2, the conserved moieties were used for moiety decomposition of the internal stoichiometric matrix, *N* ∈ ℤ^1594×2103^, corresponding to the atom mapped subset of the `iDopaNeuroC` model. The conserved moieties were used to formulate a conserved moiety fluxomic computational model based on the mathematical formulation given in Subsection 2.3.1, which enabled inference of steady state metabolic fluxes at genome-scale.

Table 1 gives the dimensions of the matrices in the mathematical formulation of conserved moiety fluxomics for the toy and `iDopaNeuroC` models. The internal reactions of the `iDopaNeuroC` model are stoichiometrically consistent, flux consistent and thermodynamically flux consistent [16]. However, a subset of the internal reactions of the `iDopaNeuroC` model were not possible to atom map, e.g., lumped reactions in lipid metabolism containing metabolites with chemically unspecified R-groups. Therefore, it is not guaranteed that every transition in the moiety transition graph can carry a net flux. Therefore in (60), the moiety balance constraint in (27) was relaxed and quadratically regularised to

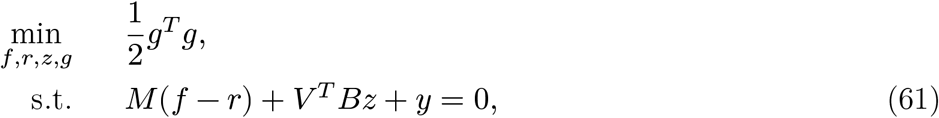

where *y* ∈ ℝ^*u*^ is a moiety balance constraint residual vector. The same relaxation and quadratic regularisation was also applied to labelled moiety balance in (28). This adaption of Problem (60) results in a strictly convex optimisation problem that always admits a feasible set and therefore an optimal solution, with a plethora of algorithms that are accompanied by mathematical guarantees of convergence to a global optimum [18].

Computation of atom mappings using Reaction Decoder Tool [29] and conserved moieties at genome-scale takes on the order of hours on a typical desktop computer, but only needs to be done once per model. Using conserved moiety fluxomics to infer a steady-state reaction flux vector at genome-scale takes ∼ 1 minute calling an industrial quality convex optimisation software (Mosek 9.2, Mosek ApS). Unless otherwise stated, all computations were implemented in the constraint-based reconstruction and analysis toolbox (COBRA Toolbox v3.4, [11]) implemented in MATLAB (Mathworks Inc.). Supplementary file 1 provides a tutorial, in the form of a MATLAB live script that implements conserved moiety fluxomics with a genome-scale model of dopaminergic neuronal metabolism and a toy model, respectively. Supplementary file 2 is the same file saved in html format (tutorial_moietyFluxomics_iDN.html).

## Discussion

In a general sense, a conserved moiety is a set of atoms whose association remains invariant with respect to the chemical transformations in a given biochemical network [20]. Previously, we mathematically defined a conserved moiety as a set of atoms, where each atom belongs to one connected component of an isomorphism class of connected components of an atom transition graph [22]. Herein we define a conserved moiety instance as a set of atoms, with the same metabolite label, where each atom belongs to one connected component of a maximal isomorphism class of connected components of an atom transition graph. Restricting all atoms in a conserved moiety instance to have the same metabolite label ensures that a conserved moiety instance is not split between multiple metabolites. A conserved moiety is a species of conserved moiety instances, where the same conserved moiety may transition between distinct metabolites so the labelling of the atoms in each instance are different, but their atomic composition is otherwise identical. In principle, this definition does not restrict each instance of a conserved moiety to be structurally identical, because rearrangement of the bonds within a conserved moiety is still possible, provided that those atoms all share the same metabolite label. Incorporation of additional information on bonds between atoms will enable future refinement of the definition of a conserved moiety (H. Rahou, unpublished).

We introduced a moiety graph decomposition of a metabolic reaction network into a set of moiety graphs. Mathematically, this enables one to relate any internal stoichiometricmatrix matrix *N* ∈ ℤ^*m*×*n*^ to the incidence matrix of a corresponding moiety transition matrix *M* ∈ {−1, 0, 1}^*u*×*v*^, where each row of *M* represents a conserved moiety instance and each column of *M* of represents the transition of a conserved moiety instance from one metabolite to another within a corresponding reaction. This decomposition is mathematically important as a stoichiometric matrix is a type of generalised incidence matrix for a hypergraph, while a moiety transition matrix is an incidence matrix for a graph. This opens up the opportunity to analyse biochemical networks using the voluminous graph theory literature, which is substantially better developed than the literature on hypergraph theory. For example, it is envisaged that this decomposition can be exploited to leverage specialised network (graph) flow algorithms, which are known to be more computationally efficient than the general linear optimisation algorithms that are currently applied to biochemical network flow problems within the constraint-based modelling field [30]. Moiety graph decomposition also avoids the artefacts associated with transformation of a biochemical hypergraph into a bipartite graph. Furthermore, it emphasises the point that biochemical networks are not hypergraphs in the general sense, but a special type of hypergraph that admits such a decomposition into a set of graphs.

We introduced conserved moiety fluxomics, which leverages moiety graph decomposition of a metabolic network to mathematically formulate a constraint-based model that enables inference of steady state metabolic fluxes that are constrained by measured isotopologue enrichment data obtained from a system at an isotopic steady state. This approach avoids combinatorial explosion of model variables for a number of reasons. In principle, the number of conserved moieties should equal the dimension of the null space of the corresponding stoichiometric matrix, that is the number of metabolites minus the rank of the stoichiometric matrix. In practice, for a genome scale model, this number is approximately equal to that ideal. It remains to refine the definition and therefore method for identification of conserved moieties so the number in practice matches the expected ideal. Nevertheless, the number of conserved moieties is always substantially less than the number of metabolites. Most reactions correspond to a small number of conserved moiety transitions and it is sufficient to only represent the flow of moieties that have the potential to be labelled. The moieties that have the potential to be labelled is known from the moiety instances present in chosen labelled substrates. Another key reason that moiety fluxomics avoids combinatorial explosion of model variables is that it is sufficient to represent constraints from those isotopologues that are actually measured. Unmeasured isotopologues would correspond to relaxed constraints so they can be omitted because mass balancing is done at the level of metabolites and reactions, with hard coupling to moiety transitions.

Conserved moiety fluxomics is dependent on the quality of the inputs, especially the metabolic network, the atom mappings and the mass isotopologue distribution data. As conserved moiety fluxomics extends an established constraint-based modelling approach it is compatible with the plethora of established constraint-based modelling methods for generating a context-specific metabolic network, given a generic metabolic model (e.g. Recon3D [9]) and context-specific data from complementary high-throughput methods (metabolomics, proteomics, transcriptomics, etc.) or low-throughput methods (e.g. manual curation of the literature). These methods have been reviewed elsewhere [31] but note that we assume the input metabolic network admits a thermodynamically feasible network flux [26] and does not force any thermodynamically infeasible flux due to erroneous combinations of reaction bounds that force flux around around a stoichiometrically balanced cycle [27]. In parallel, we have developed a novel context-specific model generation model extraction pipeline that enables one to enforce energy conservation and the second law of thermodynamics during model extraction [32]. This, in combination with the maximisation of unidirectional flux entropy ensures that inferred flux vectors do not have any component that corresponds to perpetual cycling of moieties around stoichiometrically balanced cycles. A promising future direction would be to extend conserved moiety fluxomic to incorporate thermo-chemical parameters [33] and intracellular metabolite concentrations where sampling would also enable uncertainty to be taken into account [34].

A comparison of atom mapping predictions to those obtained by manual curation of atom mappings for over five hundred reactions distributed among all top level Enzyme Commission number classes revealed that evaluated algorithms had similarly high prediction accuracy of over 90%, where an algorithm accurately predicts the atom mapping for a reaction if each atom transition for that reaction matches that obtained by manual curation. For a metabolic pathway with *n* reactions atom mapping accuracy can be approximated by *α*^*n*^ where *α* is the accuracy of an atom transition for for a single atom. Therefore, further improvement of predictive accuracy of algorithmic atom mapping as well as validation of the topology of atomically resolved metabolic networks is warranted. In principle, as conserved moiety fluxomics only involves linear constraints, it facilitates identification of either incorrect atom transitions or incorrect mass isotopologue distribution data (e.g. incorrectly identified metabolite) by identification of outlier values of mass isotopologue constraint residual variables and subsequent topological analysis of predicted versus measured labelling patterns for the corresponding conserved moieties.

The suffix -ome as used in molecular biology to mean totality with respect to the genome. Therefore, strictly speaking, fluxomics means quantification of reaction flux at genome-scale and not just for metabolism. The convex optimisation formulation underlying conserved moiety fluxomics is compatible with inference of metabolic fluxes using whole-body models of metabolism, e.g., for humans [35]. Moreover, while the focus of this paper is on genome-scale quantification of metabolic reaction flux, the mathematical approach should, in principle be applicable for any biochemical network. To enable such an approach, conserved moieties beyond metabolism would need to be computed. Here it may be useful to consider different timescales upon which a given moiety is approximately invariant. For example, this would enable definition of conserved moieties for macromolecular synthesis with timescales much longer than for metabolism [36]. More generally, the formulation of conserved moiety fluxomics should enable extension toward isotopic non-stationarity. A monolithic approach would be to define moiety fluxes for each measured time-point, set moiety balance constraints to equal the difference in moiety abundance, constrain moiety abundance with measured metabolite concentrations and add continuity constraints. An iterative approach would be to consider a sequence of convex optimisation problems, upon which there is a substantial literature regarding convergence of such iterates, e.g., criteria for well behaved perturbation to optimal solution vectors as a function of perturbed input parameters [37]. A further potential extension is the inclusion of isotopomer kinetic constraints, which are not represented in our mathematical forumulation.

## 4 Conclusions

We demonstrate how identification of the set of conserved moieties for a given metabolic network leads to a novel moiety graph decomposition of a stoichiometric matrix that can be used to linearly relate metabolic reaction fluxes to the rate at which conserved moieties transition between metabolites in a manner that avoids combinatorial explosion of model variables. A non-linear yet strictly convex objective is minimised subject to linear constraints to ensure that inferred fluxes are compatible with given isotopically labelled input and measured mass isotopologue data as well as first principles, such as mass conservation, energy conservation and the second law of thermodynamics. Numerical results demonstrate the computational tractability of conserved moiety fluxomics at genome-scale given stationary mass isotopologue distribution data from an *in vitro* dopaminergic neuronal culture fed isotopically labelled glucose. Conserved moiety fluxomics unites the strengths of constraint-based modelling and metabolic flux analysis to enable inference of steady-state metabolic reaction fluxes at genome-scale given stationary isotopic labelling data.

## Acknowledgements

RF and HSH were supported by the U.S. Department of Energy, Offices of Advanced Scientific Computing Research and the Biological and Environmental Research as part of the Scientific Discovery Through Advanced Computing program, grant #DE-SC0010429. RF, GP, AH, and TH were supported by the European Union’s Horizon 2020 research and innovation programme, for the project SysMedPD, under grant agreement No 668738. RF and TH were funded by the European Union’s Horizon Europe Framework Programme, for project Recon4IMD, under grant agreement 101080997.

## Author contributions

Ronan M.T. Fleming: Conceptualization, Methodology, Software, Validation, Formal analysis, Investigation, Writing - Original Draft, Writing - Review & Editing, Visualization, Funding acquisition

German Preciat: Investigation, Data Curation, Visualization, Writing - Review & Editing

Luojiao Huang: Investigation, Visualization, Writing - Review & Editing

Hulda S. Haraldsdottir: Formal analysis, Methodology, Writing - Review & Editing

Ines Thiele: Writing - Review & Editing

Amy Harms: Resources, Writing - Review & Editing, Supervision

Thomas Hankemeier: Resources, Writing - Review & Editing, Supervision, Funding acquisition

## A On moiety splitting of a stoichiometric matrix

Previously, we proposed an approach to split a stoichiometric matrix into the sum of a set of matrices, each corresponding to a conserved moiety [22]. We first recall this result, then relate it to the moiety decomposition proposed in (7). Given an atom transition graph 𝒢(𝒳, ℰ, ℋ) between a set of molecules 𝒱, where *m* ≔ |𝒱|, a *conserved moiety vector* 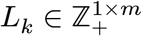 is a non-negative integer (row) vector, where *L*_*k,i*_ is the number of instances of the *k*^*th*^ conserved moiety in molecule 𝒱_*i*_. There is a one to one relationship between each conserved moiety vector and each maximal graph isomorphism class. Therefore, an atom transition graph gives rise to a set of |ℐ|conserved moiety vectors, which can be concatenated to form a *conserved moiety matrix* 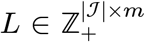,which is orthogonal to, ℛ(*N*) that is *L* · *N* = 0. Furthermore, the following matrix splitting exists

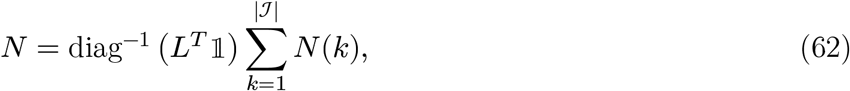

where *N*(*k*) ∈ ℤ^*m*×*n*^ is a *moiety subnetwork matrix*, given by

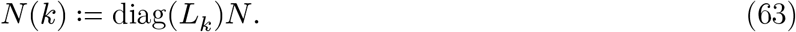

Inserting (6) into (7), one obtains the following splitting of a stoichiometric matrix

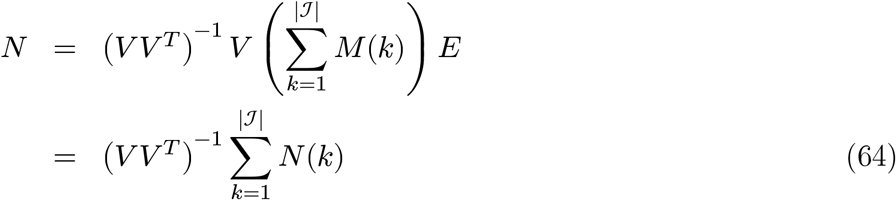

where *N*(*k*) is given by

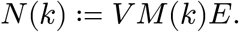

That is, the *k*^*th*^ moiety subnetwork matrix may be decomposed into the product of a matrix that maps metabolites to moieties, *V*, the incidence matrix of the *k*^*th*^ moiety subgraph, *M*(*k*), and a matrix that maps moiety transitions to reactions, *E*.

Comparing (64) and (62), we conclude that

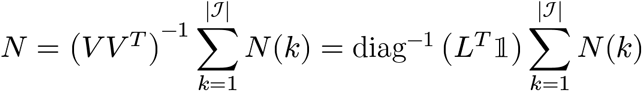

and that

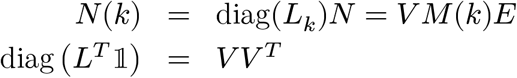

establishing a relationship between moiety splitting [22] and moiety decomposition (7) of a stoichiometric matrix.

## Footnotes

1 Huang L, Preciat G, Alarcon-Gil J, Moreno EL, Wegrzyn A, Schymanski E, Harms A, Fleming RMT, Hankemeier T, fluxTrAM: Integration of tracer-based metabolomics data into atomically-resolved genome-scale metabolic networks for metabolic flux analysis; application to human neuronal metabolism (submitted).

